# Orally Bioavailable Endochin-like Quinolone Carbonate Ester Prodrug Reduces *Toxoplasma gondii* Brain Cysts

**DOI:** 10.1101/2020.03.19.999664

**Authors:** J. Stone Doggett, Tracey Schultz, Alyssa J. Miller, Igor Bruzual, Sovitj Pou, Rolf Winter, Rozalia Dodean, Lev N. Zakharov, Aaron Nilsen, Michael K Riscoe, Vern B Carruthers

**Author notes:** Address Correspondence to J. Stone Doggett.

## Abstract

Toxoplasmosis is a potentially fatal infection for immunocompromised people and the developing fetus. Current medicines for toxoplasmosis have high rates of adverse effects that interfere with therapeutic and prophylactic regimens. Endochin-like quinolones (ELQs) are potent inhibitors of *Toxoplasma gondii* proliferation *in vitro* and in animal models of acute and latent infection. ELQ-316, in particular, was found to be effective orally against acute toxoplasmosis in mice and highly selective for the *T. gondii* cytochrome *b* over the human cytochrome *b*. Despite oral efficacy, the high crystallinity of ELQ-316 limits oral absorption, plasma concentrations and therapeutic potential. A carbonate ester prodrug of ELQ-316, ELQ-334, was created to decrease crystallinity and increase oral bioavailability, which resulted in a six-fold increase in both *C*_max_ (maximum plasma concentration) and AUC (area under the curve) of ELQ-316. The increased bioavailability of ELQ-316, when administered as ELQ-334, resulted in greater efficacy than the equivalent dose of ELQ-316 against acute toxoplasmosis and had similar efficacy against latent toxoplasmosis compared to intraperitoneal administration of ELQ-316. Carbonate ester prodrugs are a successful strategy to overcome the limited oral bioavailability of ELQs for the treatment of toxoplasmosis.

## Introduction

*Toxoplasma gondii* infection is highly prevalent in humans, parasitizing billions of people.(1) In the great majority of infections, symptoms are not appreciable; however, infected individuals are at risk of developing toxoplasmosis if they become immunologically comprised by HIV infection, cancer or immunosuppressive therapies. Infection during pregnancy can cause fetal demise, and severe congenital neurological and ocular damage. Outside of immunocompromised populations, otherwise healthy people may develop eye disease from *T. gondii* infection that can progress to blindness. Additionally, *T. gondii* causes severe infection in wild and domesticated animals and may threaten endangered species.(2, 3)

Current multidrug regimens for toxoplasmosis have had high rates of adverse events leading to discontinuation in 30% of patients.(4-6) The most common adverse events were rash, diarrhea, hematologic and hepatic toxicity.(5) High rates of hematologic toxicity are related to pyrimethamine inhibition of the host dihydrofolate reductase enzyme, and high rates of allergic reactions and overlapping toxicities with medications used in populations that are susceptible to toxoplasmosis. These groups also have higher rates of severe reactions such as toxic epidermal necrolysis and Stevens-Johnson syndrome that may be fatal.(7) The toxicities of current therapy are made worse by the prolonged exposure that patients must undergo in the initial treatment phase and the suppressive secondary prophylaxis phase. In part, the long duration of treatment is required because *T. gondii* tissue cysts are not eradicated by current regimens. New treatments that are less toxic and diminish or eliminate latent *T. gondii* tissue cysts would greatly improve outcomes for patients with toxoplasmosis.

Endochin-like quinolones (ELQ) are potent inhibitors of apicomplexan pathogens including *T. gondii, Plasmodium falciparum*, and *Babesia microti.* ELQ-316 was identified as a lead compound for toxoplasmosis based on efficacy and specificity for the *T. gondii* cytochrome *b* over the human cytochrome *b*.(8) Despite efficacy when administered orally in systemic mouse models of toxoplasmosis, malaria and babesiosis, ELQ-316 plasma and brain concentrations were not adequate to eliminate acute *T. gondii* brain infection.(9) Increasing the dose of ELQ-316 to attain adequate tissue concentrations was limited by the crystallinity of ELQs, because the bioavailability of ELQs decreases as the dose increases due to precipitation in the gastrointestinal tract at higher concentrations.(10) In order to improve the oral efficacy of ELQ-316, we created a carbonate ester prodrug of ELQ-316, ELQ-334, and determined the pharmacokinetic parameters and efficacy against acute and latent toxoplasmosis.

## Results

### Extended treatment of latent *T. gondii* brain infection with ELQ-316 and ELQ-271

Previously, ELQ-316 and ELQ-271 were shown to reduce established brain cysts when administered intraperitoneally for 15 days.(8) Based on these results, we tested a longer duration of treatment against established tissue cysts. In these experiments, CBA/J mice were infected with the ME49 *T. gondii* strain for 5 weeks prior to treatment to establish chronic brain tissue cysts. ELQs were administered to mice for 5 weeks IP at 5 mg/kg/d. This course reduced the number of brain tissue cysts rapidly within the first week with continued reduction compared to control at 5 weeks (Fig. 1A and 1B). Following treatment with ELQs, mice were given dexamethasone to evaluate the viability of the remaining cysts. Survival was prolonged in the majority of mice treated with ELQ-316; however, all mice eventually succumbed to infection indicating that viable cysts remained after treatment (Fig. 1C). Together, these findings suggest that prolonged treatment substantially reduces but does not completely eliminate viable cysts.

**FIG. 1.**
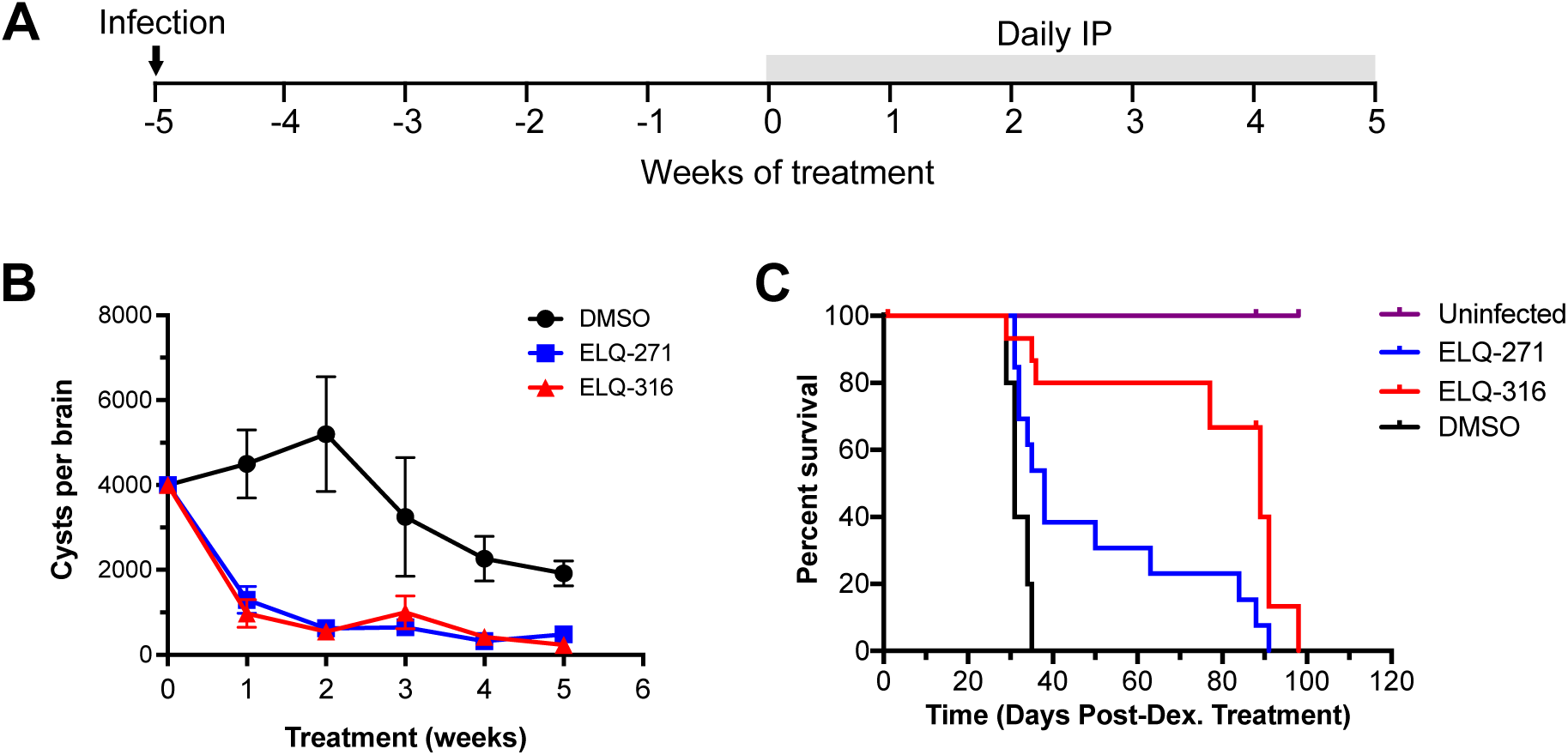
Extended treatment with ELQ-316 and 271 reduces cysts but does not eliminate viable *T. gondii* brain cysts. (A) Mice received daily treatment via IP injection for 5 weeks after being infected for 5 weeks. (B) Cysts were counted weekly in each group. Treatment with ELQ-316 and ELQ-271 decreased but did not eliminate *T. gondii* brain cysts. Cyst levels in DMSO treated mice are significantly different (p<0.05, Mann-Whitney test) from those in ELQ-271 and ELQ-316 for all time points except for week 3 between DMSO and ELQ-316. All time points and treatment groups contain cyst values from 4-8 mice except for the DMSO 4-week group, which contains data from 3 mice. (C) Mice were treated with dexamethasone following treatment with ELQs. All mice succumbed to infection. ELQ-treated mice survived longer than controls (DMSO vs. ELQ-271 p=0.017, DMSO vs ELQ-316 p=0.016; log-rank Mantel-Cox test). Uninfected, DMSO, ELQ-271, and ELQ-316 groups contained 4, 5, 13 and 9 mice, respectively. IP, intraperitoneal; PO, *per os*; Dex, dexamethasone; error bars, Standard Deviation.

### Chemical Synthesis of ELQ-334

A prodrug form of ELQ-316 was synthesized to increase oral bioavailability. ELQ-334 has been previously reported without detail of its synthesis and structural characterization.(11) In this manuscript we describe its synthesis with full characterization of the structure including X-ray diffraction analysis of ELQ-334 crystals. Following a procedure modified from Miley et al.,(12) 6-fluoro-7-methoxy-2-methyl-3-(4- (4-(trifluoromethoxy)phenoxy)phenyl)quinolin-4-yl ethyl carbonate (ELQ-334) was obtained by reacting ethyl chloroformate with 6-fluoro-7-methoxy-2-methyl-3-(4-(4- (trifluoromethoxy)phenoxy)phenyl)quinolin-4(1H)-one (ELQ-316)(13) in the presence of sodium hydride at 60°C in 95% yield as a white solid (Fig. 2). Whereas ELQ-316 decomposes at 314°C, ELQ-334 exhibits a melting point of 140°C, consistent with a significant loss of crystallinity.

**FIG. 2.**
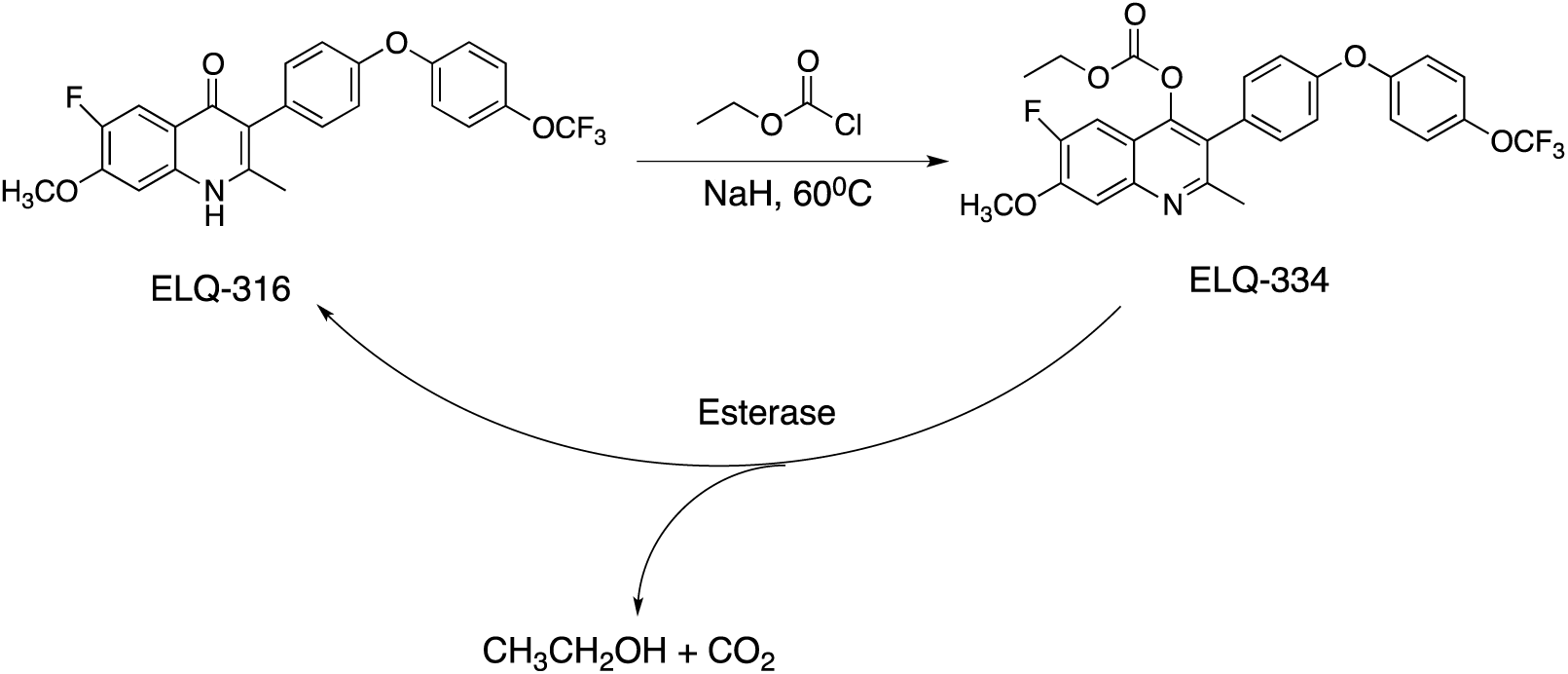
Synthesis and presumed in vivo conversion of ELQ-316 and ELQ-334.

The X-ray crystal structure of ELQ-334 contains two symmetrically independent molecules as shown in Fig. S1. The central planar quinoline ring and the first phenyl ring of the diaryl ether is twisted with a 68° torsion angle around the C2-C15 bond. Weak intermolecular hydrogen bonding networks between C-H…N and C-H…OCH3 are also observed in the crystal structures. The packing does not provide any evidence that the molecules in the crystal are connected via π–π stacking interactions. The diaryl ether group is significantly twisted from the central quinoline aromatic group to disrupt possible intermolecular π–π interactions. The absence of π–π stacking is consistent with the low melting point of 140°C.

### Noncompartmental Pharmacokinetic analysis of ELQ-316 and ELQ-334

ELQ-316 and ELQ-334 were administered orally to mice, and then plasma and brain concentrations were measured to compare the maximum concentration (*C*_max_), time to *C*_max_ (T_max_), area under the curve from time 0-96hr (AUC_0-96_), and half-life (t_1/2_) of ELQ-316 from ELQ-316 and ELQ-334. Compounds were dissolved in polyethylene glycol (PEG) 400 at a single dose of 10 mg/kg of ELQ-316. ELQ-334 yielded a plasma *C*_max_ of 4,378 ng/ml compared to 721 ng/ml of ELQ-316 from ELQ-316. The T_max_ of both compounds was 4 hours and the AUC_0-96_ was also increased approximately 6-fold to 115,195 ng/ml*h by the carbonate ester promoiety. The ELQ-316 plasma concentration of 1,665 ng/ml from ELQ-334 at 24 hours and the t_1/2_ of 11.6 hours predicts efficacy with once daily dosing. The brain tissue concentrations of ELQ-316 from ELQ-334 were 1,543 ng/ml at 4 hours and 405 ng/ml at 24 hours. The brain tissue to plasma concentration ratio was 0.35 at plasma T_max_. By comparison, the brain concentrations achieved from the oral dose of ELQ-316 were 165 ng/g at T_max_ and 75 ng/g at 24 hours. The mean plasma concentration of ELQ-334 did not exceed 238 ng/ml (5% of *C*_max_) indicating that conversion from the prodrug ELQ-334 to active compound ELQ-316 occurs very quickly and, may be caused in part by esterases present in the gastrointestinal tract. Pharmacokinetic analysis was performed with PKSolver software.(14)

### Efficacy of ELQ-334 against acute toxoplasmosis

ELQ-334 increased survival in mice at 5 mg/kg against a virulent *T. gondii* strain that is uniformly fatal in mice. ELQ-334 was given orally to mice at 1 or 5 mg/kg/d for 5 days following infection with RH strain *Toxoplasma gondii* for 24 hours. Treatment with 5 mg/kg/d prolonged survival in all mice with 2 out of 4 mice surviving through the conclusion of the experiment at 33 days. Control mice and mice treated with 1 mg/kg/d ELQ-334 were euthanized 6 and 7 days after displaying overt signs of infection, respectively. In the group treated with 5 mg/kg/d of ELQ-334, mice were euthanized at day 17 and day 20 after infection. The survival of the mice treated with ELQ-334 at 5 mg/kg was statistically greater than the control mice (p=0.008). In previously published experiments of ELQ-316 at 5 mg/kg/d in the same model, mice were euthanized at 13 days wherein systemic infection was cleared, but brain infection progressed.(15) These results suggest that the increased plasma concentration from ELQ-334 either prevents brain infection from becoming established or results in brain concentrations of ELQ-316 that are adequate to treat acute infection.

### Treatment of latent *T. gondii* brain infection with ELQ-334

ELQ-334, the carbonate ester prodrug of ELQ-316, was tested against latent *T. gondii* infection (Fig. 5). ELQ-334 given orally at 10 mg/kg for 2 weeks reduced the number of *T. gondii* cysts 67% compared to control mice that received vehicle alone and reduced the number of parasite genomes per brain 82% compared to control mice (Fig. 5B and 5C). *T. gondii* brain cysts from mice that were treated with ELQ-334 were smaller in diameter than cysts from control mice (Fig. 5D), thus providing a potential explanation for the greater reduction in parasite genomes per brain than cysts per brain due to treatment.

**FIG. 3.**
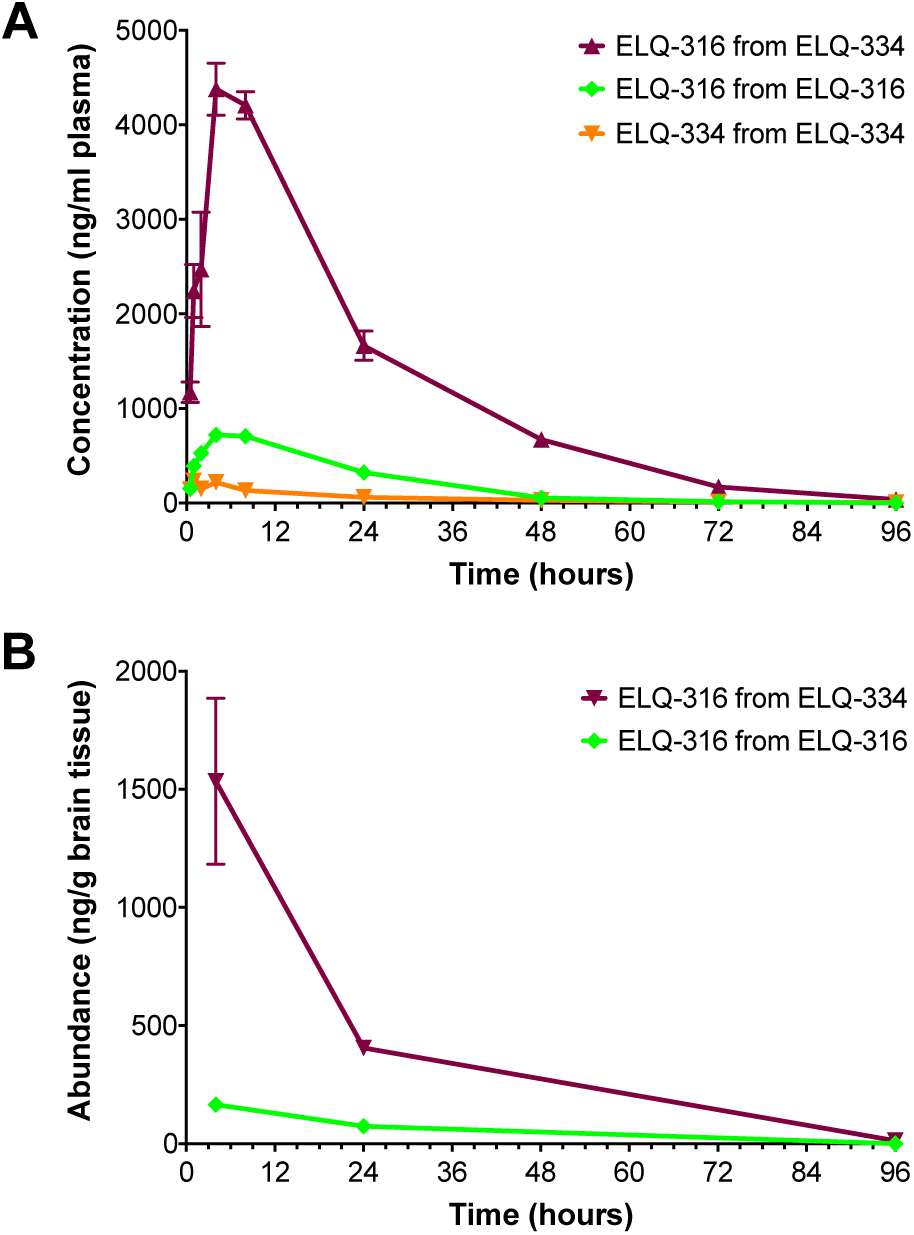
Pharmacokinetic study of ELQ-316 and ELQ-334 from oral administration of ELQ-316 or ELQ-334. Molar equivalents of 10mg/kg ELQ-316 and ELQ-334 were administered in a single dose to mice via oral gavage in PEG 400. (A) Plasma concentrations of ELQ-316 and ELQ-334 over time after administration of ELQ-316 and ELQ-334. (B) Brain tissue concentrations of ELQ-316 after administration of ELQ-316 or ELQ-334. Error bars, standard error of the mean.

**FIG. 4.**
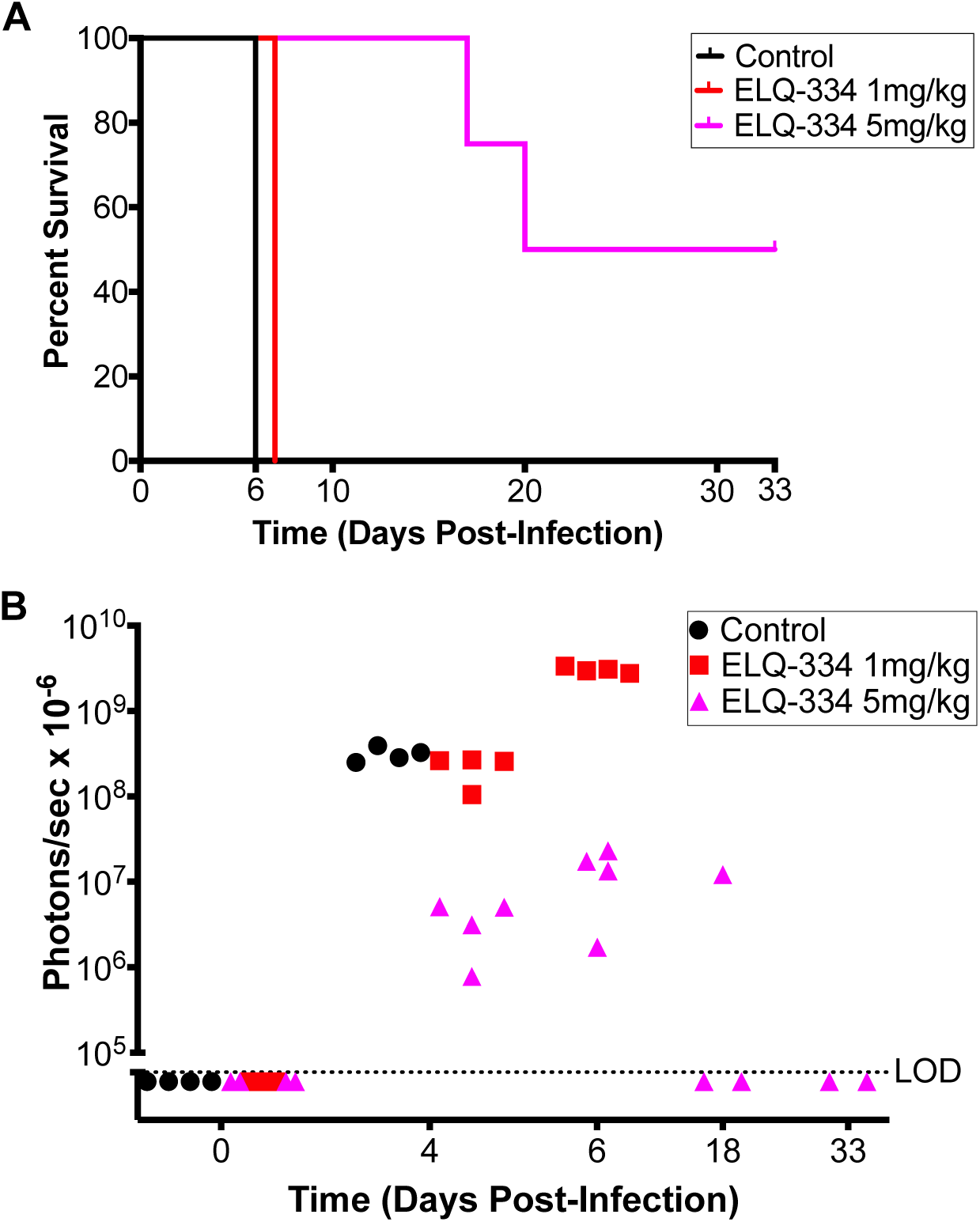
Efficacy of oral treatment with ELQ-334 against acute toxoplasmosis. Mice were infected with Type I *T. gondii* expressing firefly luciferase on day 0 followed by daily treatment for 5 days starting day 1. (A) Survival of mice treated with 5 mg/kg ELQ-334 was statistically greater than controls, p=0.008 calculated by log-rank test. (B) Luminescence in mice measured during and after treatment. LOD, limits of detection.

**FIG. 5.**
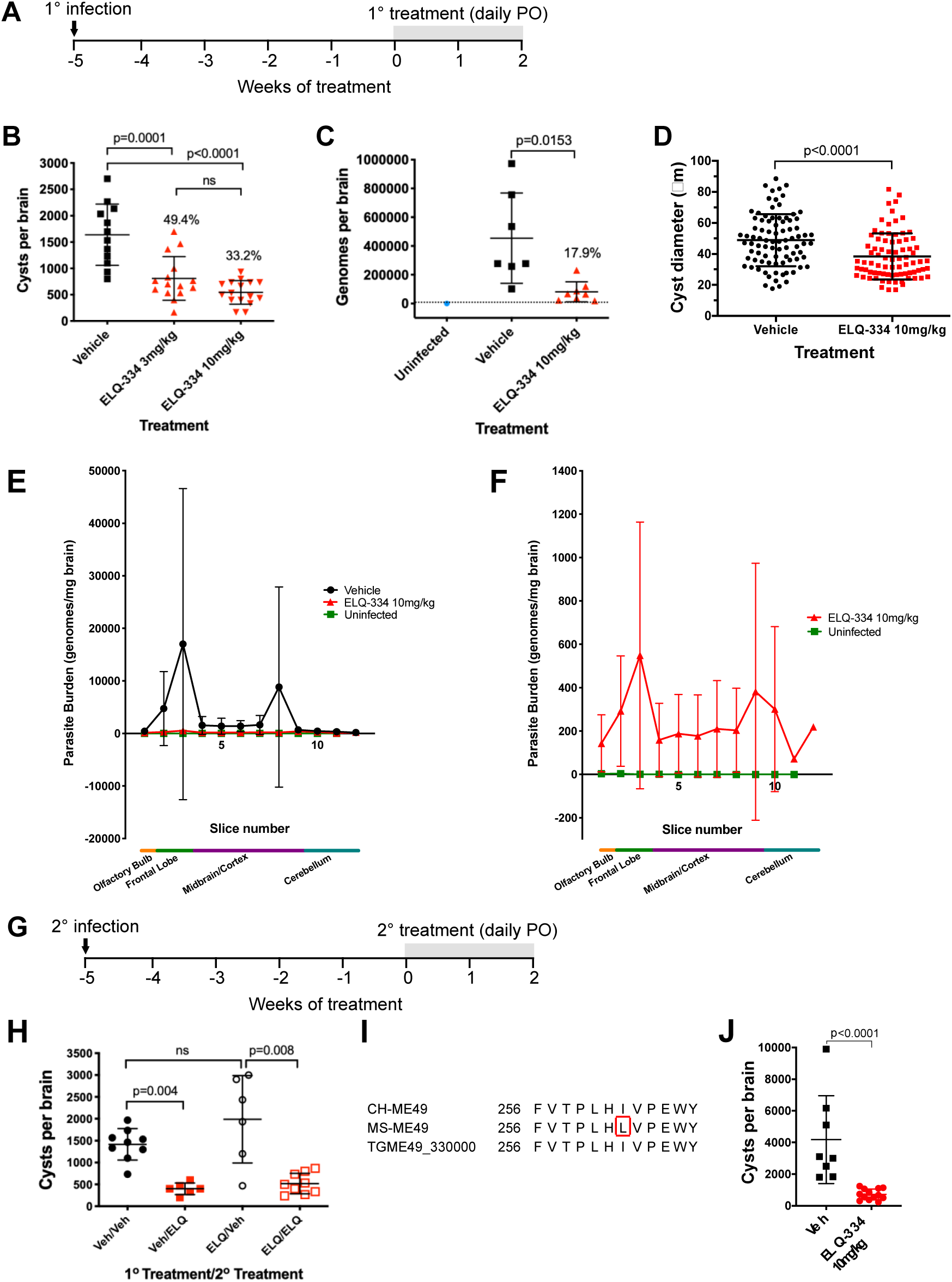
Efficacy of oral treatment with ELQ-334 against established *T. gondii* brain cysts. (A) Mice were infected for 5 weeks prior to daily treatment for 2 weeks. (B) Number of *T. gondii* brain cyst in mice treated with 3 mg/kg and 10 mg/kg of ELQ-334. Indicated p values are from an ordinary ANOVA with multiple comparisons and Tukey’s correction. One outlier was removed from the vehicle group based on ROUT analysis (Q value 0.1%). Cyst values in vehicle, ELQ-334 3 mg/kg, and ELQ-334 10 mg/kg groups are from 12, 15 and 16 mice, respectively. (C) Genomes per brain in mice treated with ELQ-334. Indicated p values are from an ordinary ANOVA with multiple comparisons and Tukey’s correction. One outlier was removed from the vehicle group based on ROUT analysis (Q value 0.1%). Data are from 7 vehicle treated mice and 8 ELQ-334 treated mice. (D) Diameter of cysts from mice treated with ELQ-334. P value is from a Mann-Whitney test. Data are from 5 brain samples per group, with 13-21 cysts measured per sample. (E) Distribution of *T. gondii* in the brains of infected mice. Data are from 8 mice per group for vehicle and ELQ-316. One mouse was used for the uninfected control. (F) Regraphing of the distribution of *T. gondii* in the brains of infected mice treated with ELQ-334 to visualize the pattern and its similarity to that of mice treated with vehicle in E. (G) Mice were infected with cysts from ELQ-334 treated mice for 5 weeks prior to retreatment with ELQ-334 for 2 weeks. (H) ELQ-334 treatment of mice infected with cysts from mice that were previously treated with ELQ-334. P values are from a Mann-Whitney test. Cyst levels in Veh/Veh, Veh/ELQ, ELQ/Veh and ELQ/ELQ groups are from 9, 6, 6, and 9 mice, respectively. (I) Comparison of cytochrome *b* Q_o_ site sequence of MS-ME49 strain to CH-ME49 strain. PO, per os; veh, vehicle, ELQ, ELQ-334; error bars, standard deviation. (J) ELQ-334 treatment of mice infected with CH-ME49. P value is from a Mann-Whitney test.

The presence of brain cysts remaining after treatment raised the possibility of inherently resistant populations of cysts or acquired drug resistance. The distribution of the cysts in the brain was examined by mechanically slicing brains into even sections from the anterior to posterior direction and estimating the total number of *T. gondii* genomes per mg of brain tissue using QRT-PCR of genomic DNA; however, no difference was observed between treated and control mice, indicating that the susceptibility of cysts to ELQ-334 was not related to cyst location (Fig. 5E and 5F). *T. gondii* exposed to ELQ-316 were tested for acquired resistance by infecting naïve mice with cysts from ELQ-334 treated mice and then treating again with ELQ-334. The number of cysts after treatment were equivalent to the original experiment in the mice that received ELQ-334 and the mice that received only vehicle (Fig. 5H).

The cytochrome *b* gene was sequenced to determine if acquired ELQ-316 resistance mutations resulted in drug resistance in the remaining tissue cysts. The MS-ME49 strain that was used in the latent toxoplasmosis experiments was found to have a single nucleotide substitution resulting in a change from isoleucine to leucine at position 262 in the Qo site of cytochrome *b* compared to CH-ME49 (a different lineage of ME49, see methods), the Type I RH strain used in the acute infection model and the cytochrome *b* sequence in the Toxodb database, TGME49_330000. This mutation has previously been associated with atovaquone resistance and is located adjacent to the highly conserved PEWY region in the Qo site (Fig. 5I).(16) The I262L substitution was discovered after the above experiments were completed. This substitution was determined to be present prior to these experiments by sequencing the cytochrome *b* gene of MS-ME49 *T. gondii* that was not exposed to ELQ-316, ELQ-334 or other compounds.

ELQ-316 resistance from cytochrome *b* Qi site mutations in *T. gondii* and *B. microti* has established the Qi site as the primary target of ELQ-316.(11, 15) The susceptibility of the CH-ME49 strain was tested to determine if the I262L mutation resulted in resistance or increased susceptibility (Fig. 5J). The remaining number of *T. gondii* cysts after treatment with ELQ-334 was not significantly different than the experiments that used the MS-ME49 strain, indicating that this substitution did not cause resistance or increased susceptibility.

## Discussion

A drug capable of eradicating *T. gondii* tissue cysts from infected individuals would prevent recurrent toxoplasmosis in people who have suffered acute infection and potentially could be used to prevent toxoplasmosis in immunocompromised individuals with evidence of latent infection. A drug that eradicates cysts may also be effective over a shorter duration than current regimens and prevent relapses of ocular toxoplasmosis that cause scarring and vision loss. Specifically, the one-year treatment of congenital toxoplasmosis and indefinite secondary prophylaxis in immunocompromised patients could be shortened to limit drug toxicity.

Over the last decade, a number of compounds with efficacy against brain tissue cysts have been identified.(8, 17-19) These compounds reduced tissue cysts but did not eliminate cysts in mouse models with intact immunity. The cause of inherent drug refractivity in latent tissue cysts has not been determined. It is also not clear if drug refractivity is specific to individual inhibitors or is a general mechanism that affects many drugs. A lack of metabolic vulnerability in non-replicating bradyzoites and decreased drug penetration through the cyst wall are among the possible explanations for drug resistance.

Extended treatment with ELQ-316 via IP injection for 5 weeks did not eradicate latent infection. Although the number of cysts was greatly reduced, the remaining cysts from the 5-week treatment were viable. *T. gondii* brain tissue cysts are dynamic over the time course of infection with cysts exhibiting varying degrees of replication and cyst packing density.(20) *T. gondii* bradyzoites express isoforms of carbon metabolism enzymes, lactate dehydrogenase and enolase, that favor glycolysis.(21-24) It has been reported that bradyzoites also lack a functional tricarboxylic acid (TCA) cycle.(25, 26) Considering these shifts in bradyzoite metabolism, cysts with replicating parasites would be vulnerable to ELQs, and cysts that are fully committed to dormancy and not reliant on oxidative phosphorylation would survive. On the other hand, cytochrome *b* inhibition induces the stage transition to bradyzoites.(27) ELQs may drive cysts with replicating parasites into latency, stopping the growth of cysts. The smaller cyst size in the ELQ treated mice may result from this growth limiting effect or be associated with an inherently recalcitrant population. That being said, the large and continued reduction in the number of cysts indicates that the great majority of brain cysts are vulnerable to cytochrome *b* inhibition.

Intermittent treatment was given both to test the efficacy of a prolonged duration and to determine if this strategy would allow cysts to become vulnerable by relieving drug pressure and allowing replication. Although the cysts were not eliminated, the number of cysts in treatment and controls continued to decline over time. The question remains whether the ultimate nadir of cyst numbers in mice was further reduced by ELQ treatment or if the cyst number in control mice would eventually reach the low numbers achieved with ELQ-316 treatment. Interestingly, previous experiments that evaluated brain cysts over 8 weeks showed an increase in the number and size of cysts.(20) Here we observed that cyst numbers began to decrease after 7 weeks in control mice and continued to decrease out to 10 weeks. In the subsequent ELQ-334 experiment, surviving cysts did not differ from controls in distribution or have acquired cytochrome *b* resistance mutations, further underscoring the inherent drug refractive properties of cysts.

An oral agent to treat toxoplasmosis is optimal for prolonged courses of therapy and in settings with limited health care resources. The carbonate ester promoiety of ELQ-334 disrupted π–π stacking leading to the enhanced oral bioavailability of ELQ-316 6-fold, resulting in a plasma concentration of 9.5 µM at *C*_max_ and 3.6 µM at 24 hours and the brain concentrations to 3.3 µM at *C*_max_ and 0.88 µM at 24 hours. These concentrations are more than 1,000-fold greater than the IC_50_ (50% inhibitory concentration) against *T. gondii*. The carbonate ester promoiety is also beneficial in that ELQ-334 was rapidly metabolized to ELQ-316, which limits the host exposure to the prodrug form. ELQ-334 treatment was more effective than ELQ-316 with 50% survival in mice compared to previous studies of orally administered ELQ-316, in which mice did not survive more than 8 days after completion of treatment.(9) Moreover, oral treatment with 10 mg/kg ELQ-334 was effective at decreasing the number of tissue cysts to a similar degree as intraperitoneal injections at 5 mg/kg. Despite not eradicating cysts, ELQ-334 quickly decreased the number of cysts that were established over 5 weeks under conditions where meningeal inflammation would be minimal and penetration through the blood brain barrier is important. The survival of mice during prolonged administration of ELQ-316 both orally and IP also demonstrates that these doses are tolerated well and cumulative toxicity does not limit efficacy in mice.

Complete elimination of tissue cysts would likely have significant clinical benefit; however, a drug that achieves this goal remains elusive. That being said, further research to understand the effect of ELQ-334 on tissue cysts and their inherent resistance may provide a means to exploit the vulnerabilities of bradyzoites in deep latency and find drug combinations that eliminate cysts. More urgently, drugs for toxoplasmosis that are more effective and better tolerated are needed. Overall, ELQ-334 is a highly promising candidate for the treatment of toxoplasmosis. Compared to atovaquone, a well-tolerated, clinically used drug, ELQ-334 is more effective, achieves higher brain concentrations, and markedly less inhibition of human cytochrome *b*.(8, 28) ELQ-316 is also more potent than the other currently used drugs pyrimethamine, sulfadiazine, clindamycin, and trimethoprim-sulfamethoxazole and does not show fetal toxicity in mice.(29) Current regimens for toxoplasmosis are significantly limited by side effects that at times are severe. Given the need for better-tolerated drugs, the efficacy of ELQ-334 described in this manuscript warrants further investigation of carbonate ester prodrugs of ELQ-316 for toxoplasmosis.

## Ethics

All animal procedures and protocols were carried out in strict accordance with the Public Health Service Policy on Humane Care and Use of Laboratory Animals and Association for the Assessment and Accreditation of Laboratory Animal Care guidelines. The University of Michigan Committee on the Use and Care of Animals (Animal Welfare Assurance A3114–01, protocol PRO00008638) and the Institutional Animal Care and Use Committee (protocol #3276) of the Portland Veterans Administration Medical Center approved the animal protocol used for this study. All efforts were made to minimize pain and suffering.

## Methods

### Experimental compounds

Unless otherwise stated all chemicals and reagents were obtained from Sigma-Aldrich Chemical Company in St. Louis, MO (USA) or Combi-Blocks in San Diego, CA and were used as received. ELQ-316 was synthesized according to Nilsen et al.(13) Melting points were obtained in the Optimelt Automated Melting point system from Stanford Research System, Sunnyvale, CA (USA). GC-MS was obtained using an Agilent Technologies 7890B gas chromatography (30 m, DBS column set at 200°C for 2 min, then at 30°C /min to 300°C with inlet temperature set at 250°C) with an Agilent Technologies 5977A mass-selective detector operating at 70 eV. Silica gel chromatography was performed using an automated flash chromatography system (Biotage Isolera One, Uppsala, Sweden). ^1^H-NMR spectra were obtained using a Bruker AMX-400 NMR spectrometer operating at 400.14 MHz. Raw NMR data were analyzed using the iNMR Spectrum Analyst software. Chemical shifts were reported in parts million units (ppm), (δ) relative to either tetramethylsilane (TMS) as internal standard or residual solvent proton (7.26 ppm for deuterated CDCl_3_). Coupling constant values were reported in hertz (Hz).

### 6-fluoro-7-methoxy-2-methyl-3-(4-(4-(trifluoromethoxy)phenoxy)phenyl)quinolin-4-yl ethyl carbonate (ELQ-334)

To a stirred suspension of ELQ-316 (3.21 g, 7.0 mmol) in dry THF (100 ml) was added a 60% mineral oil suspension of NaH (560 mg, 14.0 mmol, 2.0 Eq) and the mixture was heated at 60°C for 30 minutes. Ethyl chloroformate (1.51 g, 14.0 mmol, 2 Eq) in THF (5 ml) was added, and the reaction was heated at 60°C for 5 hours. It was then cooled to room temperature and water (10 ml) was added. The resulting mixture was filtered and separated, and the organic layer was concentrated in vacuo to give 4.1 g of a white solid. The product was purified by silica gel flash chromatography using 40% ethyl acetate in hexanes to give 3.52 g (95 % yield) of ELQ-334 as a white solid. GC-MS shows one peak with 531 M^+^ (32%), 281 (100 %). MP: 140.1-140.5 °C, ^1^H-NMR (400 MHz; CDCl_3_): δ 7.52 (d, *J* = 8.0 Hz, 1H), 7.48 (d, *J* = 11.2 Hz, 1H), 7.30-7.28 (m, 2H), 7.24-7.21 (m, 2H), 7.11-7.06 (m, 4H), 4.15 (q, *J* = 7.1 Hz, 2H), 4.04 (s, 3H), 2.53 (s, 3H), 1.22 (t, *J* = 7.1 Hz, 3H). For X-Ray analysis, ELQ-334 was re-crystalized by slow evaporation from an ethanol solution.

### X-ray crystallography

Diffraction intensities for ELQ-334 were collected at 100 K on a Rigaku XtaLAB SynergyS diffractometer using CuKα radiation, λ= 1.54184 Å. The space group was determined based on intensity statistics. Absorption correction was applied by SADABS.(30) The structure was solved by direct methods and Fourier techniques and refined on *F*^2^ using full matrix least-squares procedures. All non-H atoms were refined with anisotropic thermal parameters. H atoms in Me groups were refined in calculated positions in a rigid group model without restrictions on its rotation around C-C bonds (HFIX 138 in SHELXTL).(31) Other H atoms were found on the residual density map and refined without any restrictions with isotropic thermal parameters. In the crystal there are two symmetrical molecules. All calculations were performed using the Bruker SHELXL-2014 package.(31)

### Pharmacokinetic analysis of ELQ-316 and ELQ-334

Plasma and brain concentrations of compounds were evaluated in CF-1 mice with access to food and water ad libitum at all times. ELQ-334 and ELQ-316 were dissolved in PEG 400 to a dose of 10 mg/kg with ELQ-334 adjusted to the molar equivalency of ELQ-316. These solutions were administered orally by gavage (0.1 ml per mouse). Blood samples were collected at 0.5, 1, 2, 4, 8, 24, 48, 72 and 96 h post dose (n = 3 mice per time point), with a maximum of two samples obtained from each mouse, via tail poke. Blood was collected directly into heparinized polypropylene tubes containing a cocktail of protease inhibitor, potassium fluoride, 1 M acetic acid and EDTA to minimize the potential for *ex vivo* degradation of compounds in blood/plasma samples. All plasma samples were snap-frozen on dry ice and then stored at −80°C until analysis within 6 weeks. After the blood for the 4h, 24h and 96h time points was collected, 3 mice for each drug condition were euthanized and the brains were collected, washed with PBS and snap-frozen on dry ice and stored at −80°C. Following protein precipitation with acetonitrile (2-fold volume ratio), plasma samples were analyzed via ultraperformance liquid chromatography (UPLC)-MS (Waters Micromass Quattro Premier coupled to an Acquity UPLC device operating in positive electrospray ionization multiple-reaction monitoring mode). Analyte concentrations were determined relative to calibration curves prepared in blank mouse plasma.

### Parasite strains and passage in mice

ME49 (genotype II) strain parasites were used for all chronic infection experiments but were obtained from two different laboratories. MS-ME49 was obtained from Dr. Michael Shaw (University of Michigan) who previously received it from Dr. Alan Sher’s lab (National Institutes of Health). CH-ME49 was obtained from Dr. Christopher Hunter’s lab (University of Pennsylvania). Both versions of ME49 were maintained in Swiss Webster or CBA/J mice by serial passage at 8 to 12-week intervals. A Type I RH *T. gondii* strain expressing luciferase and green fluorescence protein was used for the acute infection model.

### Treatment of latent toxoplasmosis

Unless otherwise noted, all experiments involved seven to eight-week-old recipient CBA/J female mice infected with 18 cysts of ME49 strain parasites in brain homogenate from a 5-week infected CBA/J donor mouse. Treatment of recipient mice commenced at 5-weeks post-infection and consisted of various schemes, as described below and in the results section. Experimental compounds were administered either intraperitoneally (in 0.1 ml DMSO) or via oral gavage (in 0.1 ml PEG400). Mice were humanely euthanized 2 weeks following the final injection. The mouse brains were placed in 1 ml sterile PBS and individually minced with scissors, vortexed and homogenized by passage 3-4 times through a 22 g needle and syringe. Three 10 µl samples of each brain homogenate were placed under 24×24 mm cover slips and enumerated by phase contrast microscopy blindly i.e., without the enumerating individual having knowledge of the sample identifications.

Seventeen groups of mice were included in the course treatment experiments (Fig. 1A and B). Brain cysts in group 1 were enumerated at 5 weeks post-infection (0 weeks treatment) as a reference for the time course. Groups 2, 5, 8, 11, and 14 were treated with vehicle (DMSO) for 1-6 weeks, respectively. Groups 3, 6, 9, 12, and 15 were treated with 5 mg/kg ELQ-271 for 1-6 weeks, respectively. Groups 4, 7, 10, 13, and 16 were treated with 5 mg/kg ELQ-316 for 1-6 weeks, respectively. Treatment was administered daily via intraperitoneal injection. Groups 2-13 were euthanized one day following their last treatment for enumeration of brain cysts. Groups consisted of 5 mice each except for groups 14-16, which included 5, 13, and 10 mice, respectively because of uneven attrition due to infection. Group 17 contained 4 uninfected mice. Following 6 weeks of treatment, groups 14-17 received dexamethasone (10 mg/ml) in their drinking water to elicit immunosuppression for assessing viability of residual cysts post-treatment.

To assess the distribution of residual cysts in the brain together with testing efficacy of secondary treatment (Fig. 5), groups 1-3 consisting of 20 mice each were infected for 5 weeks and orally treated with vehicle (PEG400) or 3 mg/kg or 10 mg/kg ELQ-334 daily for 2 weeks. As uninfected negative controls, groups 4-6 (5 mice each) were treated with vehicle (PEG400) or 3 mg/kg or 10 mg/kg ELQ-334 daily for 2 weeks. Two weeks following the last treatment, mice in all groups were humanely euthanized and each brain was split into right and left hemispheres. The right hemisphere was homogenized for cyst counts, measurement of cyst size by microscopy, and for extraction of genomic DNA as described below. To assess viability of residual cysts, brain homogenates were pooled within groups and injected into naïve 7-week-old female CD1 mice (3 mice per group) at 1, 0.1, 0.01, and 0.001 brain equivalents by mixing with 0, 0.9, 0.99, or 0.999 brain equivalent homogenates from groups 4-6 i.e., uninfected, treated negative control mice, to ensure injection of equivalent brain material. CD1 mice were examined for brain cysts 5 weeks post-transfer. The left hemisphere was processed as described below. Eighteen cysts in pooled homogenates from groups 1 and 3 were also used to infect naïve CBA/J mice (10 mice per group) for secondary treatment. Five weeks post-infection these mice were orally treated with vehicle (PEG400) or 10 mg/kg ELQ-334 daily for 2 weeks. Two weeks following the last treatment, brain cysts were quantified to measure the efficacy of secondary treatment.

For comparing the sensitivity of MS-ME49 and CH-ME49 to treatment (Fig. 6), groups 1 and 2 were infected with MS-ME49 and groups 3 and 4 were infected with CH-ME49. Each group contained 15 mice. Five weeks post-infection daily oral treatment was given to groups 1 and 3 with vehicle (PEG400) and groups 2 and 4 with 10 mg/kg ELQ-334. Brain cysts were enumerated 3 weeks after the last treatment. Statistical analyses of cyst burden were performed using Mann-Whitney tests. Outliers identified with the Grubb’s test were excluded.

### Quantitative analysis of brain slices

Brains of infected and uninfected mice were dissected, weighed, flash frozen, and stored at −20°C. Tissue was then thawed and sliced into sequential 1 mm sections using a model 51425 tissue slicer (Stoelting, Wood Dale, IL). Brain slices were homogenized with 50 ul of PBS in 1.5 ml Eppendorf™ tubes using a Kontes® pellet pestle (VWR™ cat. No. KT749521-1500), followed by passaging through a 17-gauge needle attached to a 1 mL syringe. Standard samples were generated by adding *T. gondii* tachyzoites to 25 mg samples of uninfected brain tissue samples at concentrations from 0.1 to 5×10^8^ parasites/mg tissue. Whole genomic DNA from samples and standards was extracted and purified using the DNAeasy Blood and Tissue Kit by Qiagen™ (cat. no. 69504) according to manufacturer’s instructions. 50 ng of genomic DNA was used per sample to run qPCR using primers against a *Toxoplasma*-specific 529 bp repeat element (Tox-9 Forward, 5′ AGGAGAGATATCAGGACTGTAG; Tox-11 Reverse, 5′ GCGTCGTCTCGTCTAGATCG). All samples were quantified by qPCR using the Bio-Rad SYBR Green master mix reagent (ThermoFisher Scientific™ cat. no. 4309155) on an Applied Biosystems Step One Plus qPCR instrument. Treated and control sample threshold cycle (CT) values were compared against the standard curve to extrapolate the number of *T. gondii* genomes per brain slice. Samples with a CT value above 40 were considered to have no detectable *T. gondii* genomes present.

### Treatment of acute toxoplasmosis

CF-1 mice that were 4-6 weeks old were inoculated intraperitoneally with 10,000 virulent Type I RH *T. gondii* tachyzoites that express firefly luciferase and GFP. After 24 hours, compounds were dissolved and administered in PEG400 via oral gavage daily for five days. Vehicle only control groups and treatment groups consisted of 4 mice per group. Mice were monitored for signs of infection, and underwent bioluminescence imaging on day 4, 6, 13 and 29. Mice were injected IP with a dose of 0.1 ml of D-luciferin (150 mg substrate/kg of body weight) dissolved in PBS. Three minutes after luciferin injection, mice were anesthetized using inhaled isoflurane and positioned ventral side up on a heated platform. Bioluminescent images were obtained using an IVIS Spectrum CT and processed using Living Image software (Perkin Elmer). Mice that developed signs of severe infection, such as >10% weight loss, lethargy, or lack of self-grooming, or at 33 days, were humanely euthanized. Analysis of survival and differences of the tissue burden of *T. gondii* infection were performed using a logrank test and unpaired t-test, respectively. GraphPad Prism 7.0 software was used for statistical analysis.

### Cytochrome *b* gene sequencing

DNA was isolated from 6 samples of MS-ME49 *T. gondii* cysts, CH-ME49 cysts and RH strain tachyzoites using the DNeasy Blood % Tissue Purification Kit (Qiagen). The cytochrome *b* coding sequences were amplified from genomic DNA and cDNA by PCR with primers 5’ATGGTTTCGAGAACACTCAGT, 3’GTATAAGCATAGAACCAATCCGGT and Phusion DNA polymerase yielding a single PCR product visualized on an agarose gel. Control PCR reactions without DNA did not yield PCR products. Amplicons were sequenced using sequencing primers 5’CTACCATGGGGACAAATGAGTTTCTGGGGTGCTACAGT and 3’ACCATTCTGGTACGATATGAAGTGGTGTTAC. Protein alignment was performed with MUSCLE (Multiple Sequence Comparison by Log-Expectation).

## Acknowledgements

This work was supported by Career Development Award BX002440 and VA Merit Review Award BX004522 to J.S.D. from the U.S. Department of Veterans Affairs Biomedical Laboratory Research and Development. We also acknowledge support for M.K.R. from NIH R01 AI100569, Peer Reviewed Medical Research Program Project PR130649, and VA Merit Review Funds from the U.S. Department of Veterans Affairs BX003312. The work was also supported by a grant from the Stanley Medical Research Institute to V.B.C.

---

**FIG S1.**
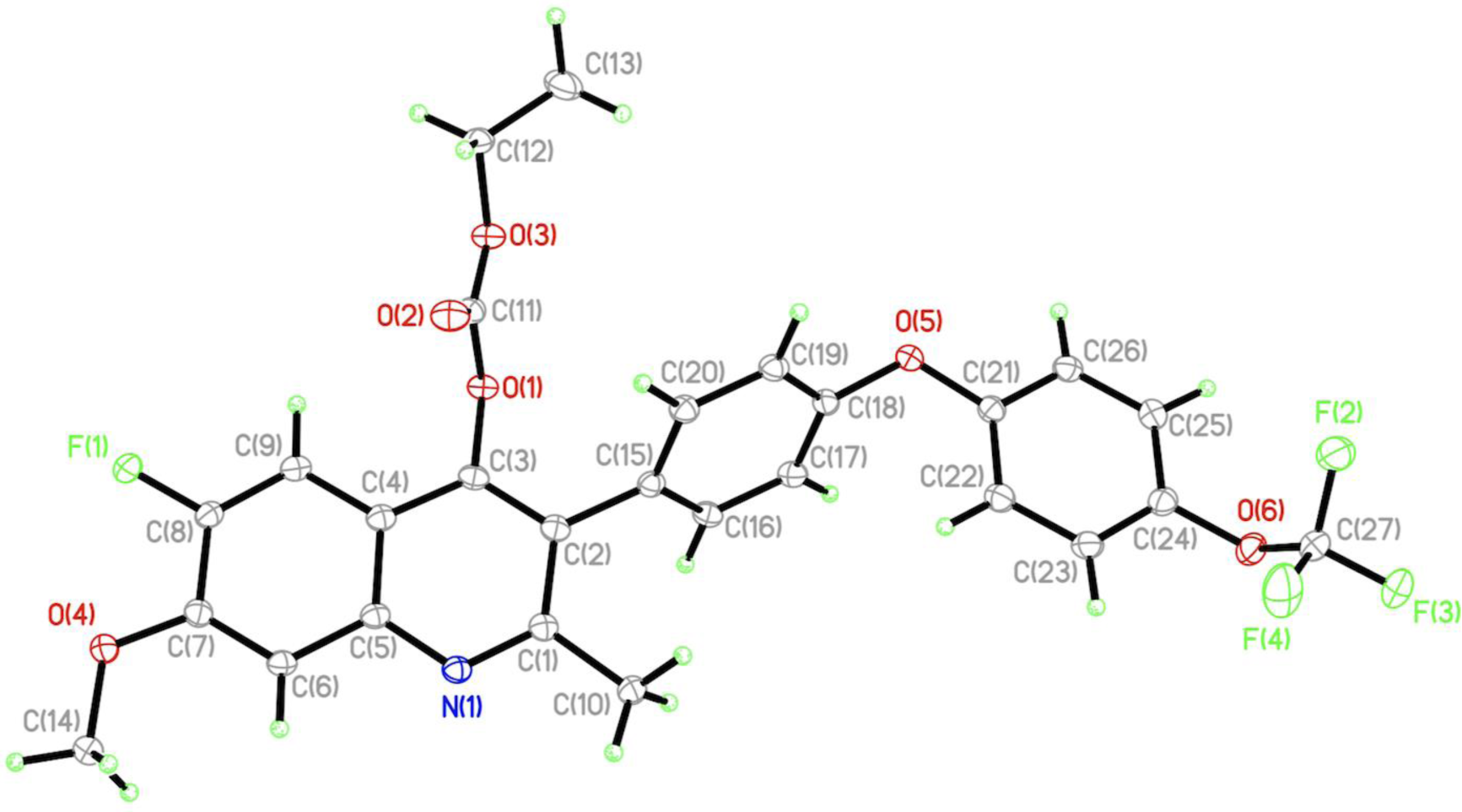
Oak Ridge Thermal Ellipsoid Plot (ORTEP) of ELQ-334

**FIG S2.**
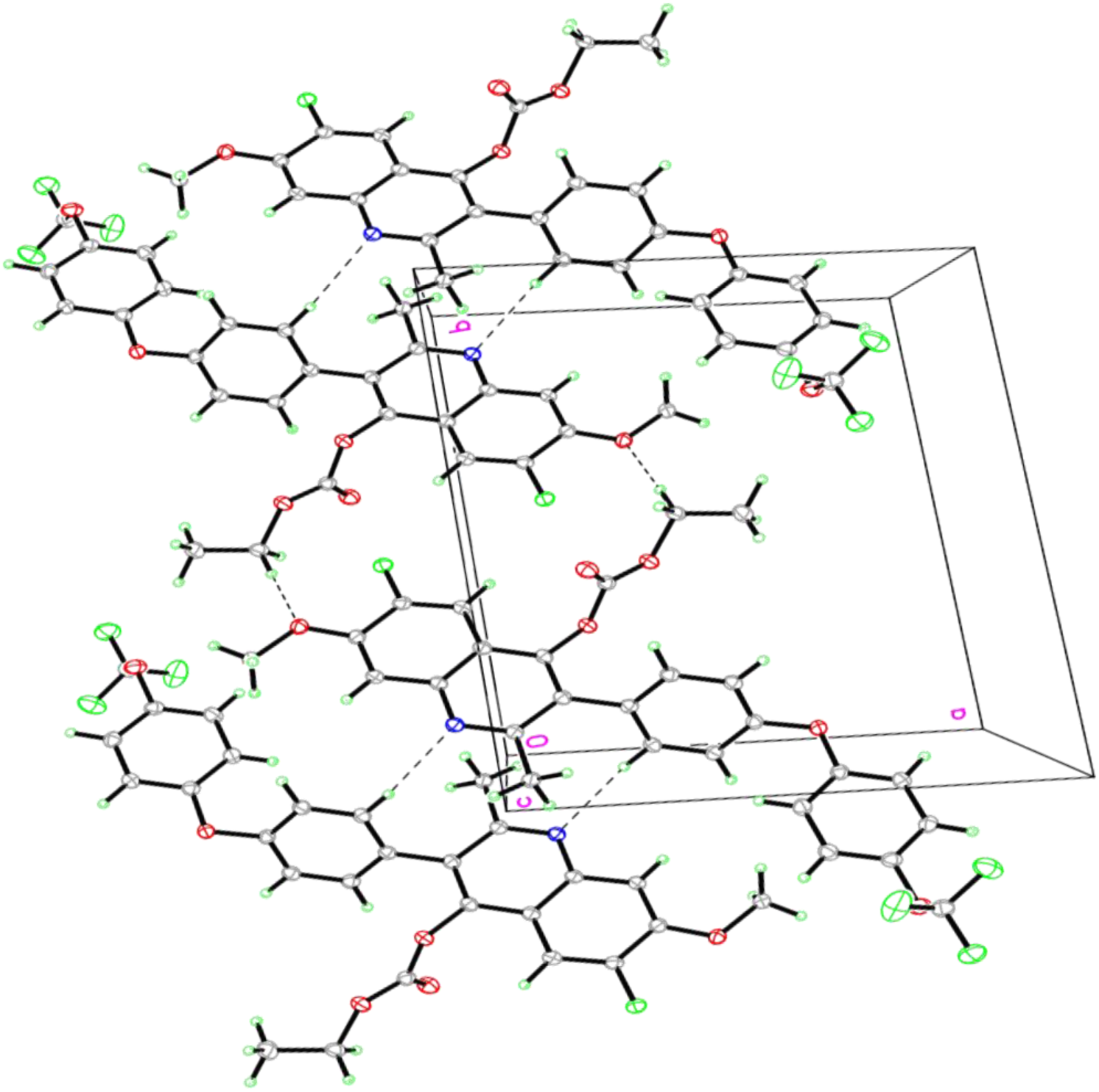
A fragment of the crystal structure of ELQ-334. Hydrogen bonds are shown by dash lines. Thermal ellipsoids are drawn at the 30% probability level.

## References

1. Tenter AM, Heckeroth AR, Weiss LM. 2000. Toxoplasma gondii: from animals to humans. Int J Parasitol 30:1217–58.

2. Jones JL, Parise ME, Fiore AE. 2014. Neglected parasitic infections in the United States: toxoplasmosis. Am J Trop Med Hyg 90:794–9.

3. Shapiro K, VanWormer E, Packham A, Dodd E, Conrad PA, Miller M. 2019. Type X strains of Toxoplasma gondii are virulent for southern sea otters (Enhydra lutris nereis) and present in felids from nearby watersheds. Proc Biol Sci 286:20191334.

4. Yan J, Huang B, Liu G, Wu B, Huang S, Zheng H, Shen J, Lun ZR, Wang Y, Lu F. 2013. Meta-analysis of prevention and treatment of toxoplasmic encephalitis in HIV-infected patients. Acta Trop 127:236–44.

5. Katlama C, De Wit S, O’Doherty E, Van Glabeke M, Clumeck N. 1996. Pyrimethamine-clindamycin vs. pyrimethamine-sulfadiazine as acute and long-term therapy for toxoplasmic encephalitis in patients with AIDS. Clin Infect Dis 22:268–75.

6. Porter SB, Sande MA. 1992. Toxoplasmosis of the central nervous system in the acquired immunodeficiency syndrome. N Engl J Med 327:1643–8.

7. Lin D, Tucker MJ, Rieder MJ. 2006. Increased adverse drug reactions to antimicrobials and anticonvulsants in patients with HIV infection. Ann Pharmacother 40:1594–601.

8. Doggett JS, Nilsen A, Forquer I, Wegmann KW, Jones-Brando L, Yolken RH, Bordon C, Charman SA, Katneni K, Schultz T, Burrows JN, Hinrichs DJ, Meunier B, Carruthers VB, Riscoe MK. 2012. Endochin-like quinolones are highly efficacious against acute and latent experimental toxoplasmosis. Proc Natl Acad Sci U S A 109:15936–41.

9. McConnell EV, Bruzual I, Pou S, Winter R, Dodean RA, Smilkstein MJ, Krollenbrock A, Nilsen A, Zakharov LN, Riscoe MK, Doggett JS. 2018. Targeted Structure-Activity Analysis of Endochin-like Quinolones Reveals Potent Qi and Qo Site Inhibitors of Toxoplasma gondii and Plasmodium falciparum Cytochrome bc1 and Identifies ELQ-400 as a Remarkably Effective Compound against Acute Experimental Toxoplasmosis. ACS Infect Dis.

10. Nilsen A, LaCrue AN, White KL, Forquer IP, Cross RM, Marfurt J, Mather MW, Delves MJ, Shackleford DM, Saenz FE, Morrisey JM, Steuten J, Mutka T, Li Y, Wirjanata G, Ryan E, Duffy S, Kelly JX, Sebayang BF, Zeeman AM, Noviyanti R, Sinden RE, Kocken CH, Price RN, Avery VM, Angulo-Barturen I, Jimenez-Diaz MB, Ferrer S, Herreros E, Sanz LM, Gamo FJ, Bathurst I, Burrows JN, Siegl P, Guy RK, Winter RW, Vaidya AB, Charman SA, Kyle DE, Manetsch R, Riscoe MK. 2013. Quinolone-3-diarylethers: a new class of antimalarial drug. Sci Transl Med 5:177ra37.

11. Lawres LA, Garg A, Kumar V, Bruzual I, Forquer IP, Renard I, Virji AZ, Boulard P, Rodriguez EX, Allen AJ, Pou S, Wegmann KW, Winter RW, Nilsen A, Mao J, Preston DA, Belperron AA, Bockenstedt LK, Hinrichs DJ, Riscoe MK, Doggett JS, Ben Mamoun C. 2016. Radical cure of experimental babesiosis in immunodeficient mice using a combination of an endochin-like quinolone and atovaquone. J Exp Med.

12. Miley GP, Pou S, Winter R, Nilsen A, Li Y, Kelly JX, Stickles AM, Mather MW, Forquer IP, Pershing AM, White K, Shackleford D, Saunders J, Chen G, Ting LM, Kim K, Zakharov LN, Donini C, Burrows JN, Vaidya AB, Charman SA, Riscoe MK. 2015. ELQ-300 prodrugs for enhanced delivery and single-dose cure of malaria. Antimicrob Agents Chemother 59:5555–60.

13. Nilsen A, Miley GP, Forquer IP, Mather MW, Katneni K, Li Y, Pou S, Pershing AM, Stickles AM, Ryan E, Kelly JX, Doggett JS, White KL, Hinrichs DJ, Winter RW, Charman SA, Zakharov LN, Bathurst I, Burrows JN, Vaidya AB, Riscoe MK. 2014. Discovery, synthesis, and optimization of antimalarial 4(1H)-quinolone-3-diarylethers. J Med Chem 57:3818–34.

14. Zhang Y, Huo M, Zhou J, Xie S. 2010. PKSolver: An add-in program for pharmacokinetic and pharmacodynamic data analysis in Microsoft Excel. Comput Methods Programs Biomed 99:306–14.

15. Alday PH, Bruzual I, Nilsen A, Pou S, Winter R, Ben Mamoun C, Riscoe MK, Doggett JS. 2017. Genetic Evidence for Cytochrome b Qi Site Inhibition by 4(1H)-Quinolone-3-Diarylethers and Antimycin in Toxoplasma gondii. Antimicrob Agents Chemother 61.

16. McFadden DC, Tomavo S, Berry EA, Boothroyd JC. 2000. Characterization of cytochrome b from Toxoplasma gondii and Q(o) domain mutations as a mechanism of atovaquone-resistance. Mol Biochem Parasitol 108:1–12.

17. Vidadala RS, Rivas KL, Ojo KK, Hulverson MA, Zambriski JA, Bruzual I, Schultz TL, Huang W, Zhang Z, Scheele S, DeRocher AE, Choi R, Barrett LK, Siddaramaiah LK, Hol WG, Fan E, Merritt EA, Parsons M, Freiberg G, Marsh K, Kempf DJ, Carruthers VB, Isoherranen N, Doggett JS, Van Voorhis WC, Maly DJ. 2016. Development of an Orally Available and Central Nervous System (CNS) Penetrant Toxoplasma gondii Calcium-Dependent Protein Kinase 1 (TgCDPK1) Inhibitor with Minimal Human Ether-a-go-go-Related Gene (hERG) Activity for the Treatment of Toxoplasmosis. J Med Chem 59:6531–46.

18. Rutaganira FU, Barks J, Dhason MS, Wang Q, Lopez MS, Long S, Radke JB, Jones NG, Maddirala AR, Janetka JW, El Bakkouri M, Hui R, Shokat KM, Sibley LD. 2017. Inhibition of Calcium Dependent Protein Kinase 1 (CDPK1) by Pyrazolopyrimidine Analogs Decreases Establishment and Reoccurrence of Central Nervous System Disease by Toxoplasma gondii. J Med Chem.

19. Benmerzouga I, Checkley LA, Ferdig MT, Arrizabalaga G, Wek RC, Sullivan WJ, Jr. 2015. Guanabenz repurposed as an antiparasitic with activity against acute and latent toxoplasmosis. Antimicrob Agents Chemother 59:6939–45.

20. Watts E, Zhao Y, Dhara A, Eller B, Patwardhan A, Sinai AP. 2015. Novel Approaches Reveal that Toxoplasma gondii Bradyzoites within Tissue Cysts Are Dynamic and Replicating Entities In Vivo. MBio 6:e01155–15.

21. Al-Anouti F, Tomavo S, Parmley S, Ananvoranich S. 2004. The expression of lactate dehydrogenase is important for the cell cycle of Toxoplasma gondii. J Biol Chem 279:52300–11.

22. Dzierszinski F, Mortuaire M, Dendouga N, Popescu O, Tomavo S. 2001. Differential expression of two plant-like enolases with distinct enzymatic and antigenic properties during stage conversion of the protozoan parasite Toxoplasma gondii. J Mol Biol 309:1017–27.

23. Ferguson DJ, Parmley SF, Tomavo S. 2002. Evidence for nuclear localisation of two stage-specific isoenzymes of enolase in Toxoplasma gondii correlates with active parasite replication. Int J Parasitol 32:1399–410.

24. Lunghi M, Galizi R, Magini A, Carruthers VB, Di Cristina M. 2015. Expression of the glycolytic enzymes enolase and lactate dehydrogenase during the early phase of Toxoplasma differentiation is regulated by an intron retention mechanism. Mol Microbiol 96:1159–75.

25. Sullivan WJ, Jr., Jeffers V. 2012. Mechanisms of Toxoplasma gondii persistence and latency. FEMS Microbiol Rev 36:717–33.

26. Denton H, Roberts CW, Alexander J, Thong KW, Coombs GH. 1996. Enzymes of energy metabolism in the bradyzoites and tachyzoites of Toxoplasma gondii. FEMS Microbiol Lett 137:103–8.

27. Bohne W, Heesemann J, Gross U. 1994. Reduced replication of Toxoplasma gondii is necessary for induction of bradyzoite-specific antigens: a possible role for nitric oxide in triggering stage conversion. Infect Immun 62:1761–7.

28. GlaxoSmithKline Research Triangle Park N. 2019. tMepron (atovaquone oral suspension) [package insert].

29. Anghel N, Balmer V, Muller J, Winzer P, Aguado-Martinez A, Roozbehani M, Pou S, Nilsen A, Riscoe M, Doggett JS, Hemphill A. 2018. Endochin-Like Quinolones Exhibit Promising Efficacy Against Neospora Caninum in vitro and in Experimentally Infected Pregnant Mice. Front Vet Sci 5:285.

30. Sheldrick GM. 1998. Bruker/Siemens Area Detector Absorption Correction Program, Madison, WI.

31. Sheldrick GM. 2015. Crystal structure refinement with SHELXL. Acta Crystallogr C Struct Chem 71:3–8.

